# EndoNB: A general strategy to study the internalization of cell surface proteins

**DOI:** 10.1101/2025.06.08.658482

**Authors:** An-Sofie Lenaerts, Jekaterina Kristal, Tai Arima, Neza Leskovar, Narjes Zeinoddin, Leonardo Almeida-Souza

## Abstract

The cell surface harbors thousands of distinct proteins whose function depend on continuous cycles of internalization and replenishment. Disturbances in this turnover are typical hallmarks of many diseases. Yet, tools to study the dynamics of most surface proteins are suboptimal or unavailable. Here, we present a new method that enables the analysis of surface protein turnover of virtually any surface protein at endogenous levels. Our approach combines CRISPR/cas9-mediated genome engineering with a cleavable recombinant probe, which addresses many of the shortcomings of current methodologies. We demonstrate the capabilities our method by studying the internalization behavior of a previously uncharacterized surface protein and by assessing the effect of ligands and activity modulators in the endocytic behavior of established receptors. In summary, our method represents a versatile strategy to explore surface protein biology and enhances our ability to study the mechanisms of membrane protein retrieval and recycling.

## Introduction

The plasma membrane is a complex cellular structure that, alongside structural lipids and sugars, harbors a vast array of distinct proteins, which represent approximately 15% of the human genome^1,2^. These proteins perform various functions such as cell protection, adhesion, migration, nutrient uptake, signaling, and cell-cell communication. Their distribution and concentration on the cell surface are tightly controlled by rapid cycles of endocytosis and exocytosis. Alterations in the dynamics of plasma membrane components have been linked to various diseases, including cancer, metabolic disorders, psychiatric conditions, and neurodegeneration^3^. Despite their importance, tools to measure the dynamics of these proteins are not widely available.

A key challenge to study the internalization of plasma membrane proteins is the requirement of a fluorescently labelled ligand or antibody against the protein of interest. Unfortunately, only a few ligands can be labelled and specific antibodies targeting the extracellular regions of membrane proteins are rare. Moreover, the bivalent nature of antibodies may lead to surface proteins clustering and/or activation, thereby interfering with their activation state, endocytic route and function^4^.

At the end of any internalization assay, there is a crucial step where the ligands or antibodies bound to non-internalized surface proteins are removed. This ensures that all the fluorescent signal derives from endocytosed surface proteins. For many of these proteins, only a fraction is internalized at a given time. Therefore, the surface signal removal is crucial for obtaining meaningful results. A few methods to remove or reduce surface signal have been used. The most widely used protocol for this step is an “acid wash”, where cells are incubated with a low pH buffer aiming the removal of antibodies^5,6^ or labelled ligands^7,8^ from the cell surface. Another commonly used protocol is fluorescence quenching, which uses anti-fluorophore antibodies^9,10^ which efficiently (but not completely) block the signal of fluorophores on antibodies or ligands^11^. A less common method uses proteases to digest surface proteins and antibodies from the cell’s surface^12,13^. A common feature of these methods is their high variability between cells, antibodies, ligands and operator and generally require extensive optimization to balance cell integrity and optimal surface signal removal.

The last two decades have seen an explosion in the number of new endocytic pathways or new adaptor proteins for well-established ones^14^. Despite these advances, we are only scratching the surface on understanding the full complexity of plasma membrane dynamics. Most endocytic pathways are defined by the behavior of only a handful of surface proteins. Given that a typical cell line harbors around 1000 different surface proteins at any given time^2^, tools allowing us to study this untapped complexity are likely to reveal novel internalization and recycling routes and significantly increase our understanding of routes we already know.

Here, we present a strategy capable of studying the internalization dynamics of potentially any membrane-resident protein at endogenous levels with an inbuilt feature that greatly simplifies the surface signal removal step. We call this strategy EndoNB (<u>Endo</u>cytosis with the aid of peptide-recognizing <u>N</u>ano<u>B</u>odies).

## Results

### The principle of EndoNB

We reasoned that an easy, biologically significant and generalizable internalization assay should have three features: i) work on endogenous proteins with minimal impact on protein function; ii) have a simple and efficient way to remove the surface signal and; iii) have easy and flexible readout methods. We solved the first feature using CRISPR/Cas9 to insert a short, 15 amino acid Alfa-tag^15^ to regions of surface proteins that do not interfere with their function or lie on flexible, disordered loops away from dimerization interfaces or ligand binding sites (when available) (Fig. 1a). AlphaFold^16,17^ was used to predict the fold of the tagged proteins and to compare it to their untagged versions. The second feature was solved by producing a recombinant protein probe consisting of a nanobody Alfa (which recognizes the Alfa-tag) and a SNAPtag (which can be fluorescently labelled) connected by a linker with a 3C protease site (Fig. 1b). The 3C protease shows good activity on ice, allowing the easy removal of the fluorescent moiety (SNAPtag) of the EndoNB probe at conditions of stalled endocytic trafficking (i.e. low temperature). With the surface signal removed, cells can be processed for microscopy or flow cytometry for easy internalization quantification (Fig. 1c), thereby achieving the third feature of our assay.

**Figure 1.**
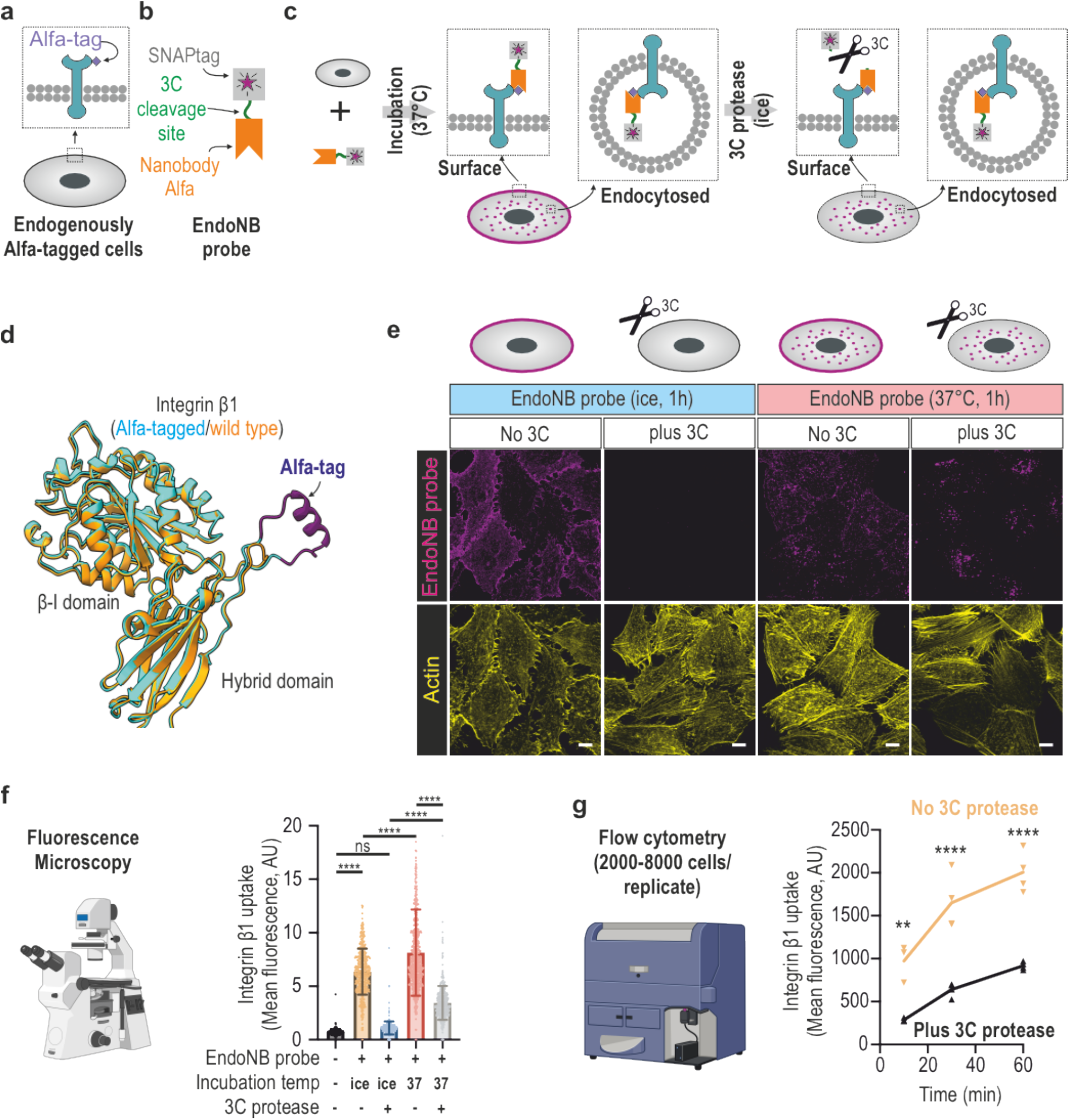
EndoNB proof-of-principle. **a-b**, Schematics of EndoNB components. **a**. Generic cell with a surface protein endogenously tagged with an extracellular Alfa-tag. **b**. Recombinant EndoNB probe: composed of a nanobody Alfa and a SNAPtag linked by a HRV 3C cleavage site. **c**, EndoNB assay principle. Cells with an endogenously Alfa-tagged surface protein are incubated with the EndoNB probe for a defined time. In this period, EndoNB will bind to the Alfa tag on the targeted surface protein and mirror the behavior of the surface protein (i.e. some EndoNB molecules are internalized into vesicles, while others will remain on non-internalized surface proteins). After incubation, cells are placed on ice to stop trafficking and a treatment with 3C protease cleaves off the fluorescent moiety of EndoNB (SNAPtag) thereby removing all remaining signal on the cell surface. **d**, Structural alignment of the hybrid and β-I domains of wild-type and Alfa-tagged Integrin β1. Both structures were generated with AlphaFold 3. **e**, Proof-of-principle assay for EndoNB using Alfa-tagged Integrin β1 cells. Images of cells incubated with EndoNB and treated or not with 3C as indicated. Images are maximum intensity projections of confocal images. Scale bar = 10 µm. **f**, Quantification of integrin β1 uptake by microscopy using EndoNB (n= 412, 461, 467, 667, 512 cells). **g**, Quantification of integrin β1 uptake by flow cytometry using EndoNB (n =4; each n represents the median fluorescence of 2000-8000 cells). The microscope and the flow cytometer illustrations were generated with Biorender. ns = non-significant, ** p> 0.01, *** p> 0.001, **** p> 0.0001. ANOVA with Tukey’s post hoc analysis.

For a proof-of-principle experiment, we used integrin β1. This integrin is expressed at high levels in many cell types and plays crucial function in cell migration during development and metastasis^18^. We endogenously Alfa-tagged this gene (ITGB1) at the codon for glycine 101, located in a flexible loop which was previously used to insert a much larger GFP without significant functional interference^19^ (Fig. 1d). To show the feasibility of our strategy, Alfa-tagged integrin β1 cells were incubated with the EndoNB probe for 1h at 37°C or on ice (i.e. no endocytosis), followed by a 1h treatment with the 3C protease on ice. We also included a control condition without the 3C protease (PBS only). As shown in figure 1e, the 3C protease treatment was very effective in removing the surface signal, bringing it to background levels in cells incubated with EndoNB on ice. Moreover, in the 1h internalization condition, the 3C protease treatment resulted in a clean vesicular signal coming only from internalized integrins (Fig. 1e). The consistent reduction in background after 3C protease treatment allowed us to easily quantify internalization using both microscopy (Fig. 1f) and flow cytometry (Fig. 1g). Thus, these proof-of-principle experiments demonstrate that EndoNB allows easy and reproducible measurements of surface protein uptake. The suitability of our approach to multiple readout modalities allows the observation of endosomal distribution of cargoes via imaging or to easily measure cargo uptake of thousands of cells in many experimental conditions using flow cytometry.

### Applying EndoNB to study the effects of ligands and functional antibodies

The working principle of EndoNB allows for the easy comparison of the endocytic behavior of receptors in the presence or absence of their cognate ligands or allosteric modulators. To test this usage, we selected three different receptors with distinct behaviors and available functional modulators: Integrin β1; the transferrin receptor, TFR; and the tyrosine receptor kinase AXL.

Over the years, a large collection of anti-integrin β1 antibodies have been developed and characterised^20^. Some of these display integrin β1 activation or inactivation activities and have been used extensively to modulate integrin β1 activity and to study specific integrin β1 subpopulations. To illustrate how EndoNB can address new questions in integrin biology, we used our assay to understand how integrin β1 -activating (12G10) and -inactivating (Mab13) antibodies affect the endocytosis of the total integrin β1 pool (Fig. 2a,b). As shown in figure 2c, Mab13 treatment led to a twofold increase in total integrin β1 uptake. On the other hand, treatments with 12G10 or a control anti-integrin αVβ3 antibody did not change total integrin β1 uptake when compared to the non-treated control. To our knowledge, this is the first demonstration of the dramatic effect of Mab13 treatment on the total endocytosis of integrin β1. As Mab13 stabilizes the inactive, closed conformation of integrin β1^21^, our results suggest that the inactive form of this integrin is biased towards its internalization. This possibility is challenged by the fact that most of the integrin β1 in the cell surface (around 80%) is reported to be in the inactive conformation^9^. However, integrin activation states fluctuate between intermediate states^22,23^ and one could argue that Mab13 increases the time integrins stay in their most endocytic-prone state.

**Figure 2.**
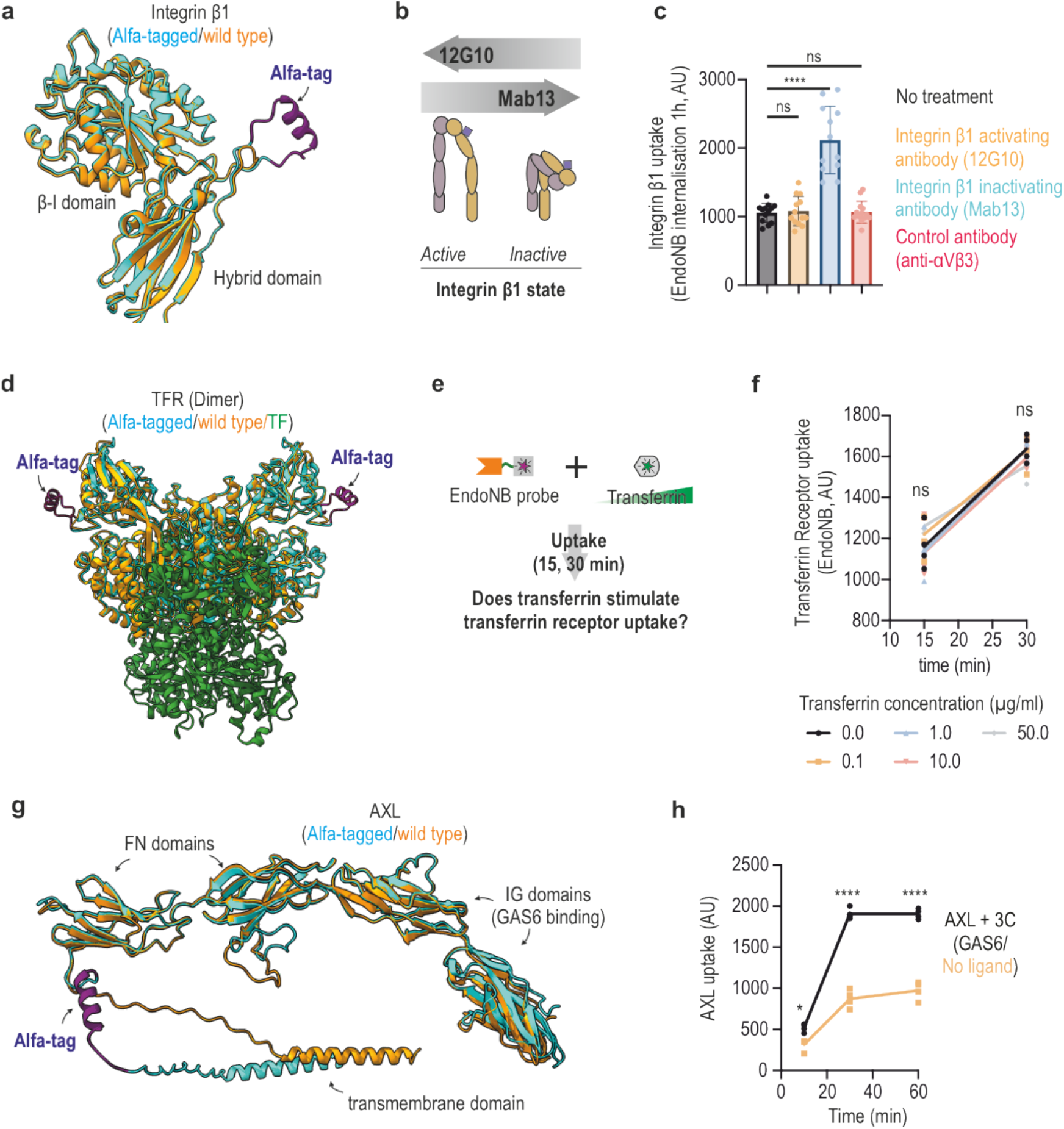
EndoNB to study the effect of ligands and functional antibodies. **a**, Structural alignment of the hybrid and β-I domains of wild-type and Alfa-tagged Integrin β1. **b**, Schematics showing the effect of 12G10 and Mab13 antibodies on integrin β1 activation state. These antibodies act by stabilizing the integrins in their active or inactive state. **c**, Quantification of integrin β1 uptake by flow cytometry using EndoNB in the presence of no antibodies (no treatment), or in the presence of 12G10 (activating), Mab13 (inactivating) or unrelated (anti integrin αvβ3 antibody). (n = 12, each n represents the median fluorescence of 2000-8000 cells). **d**, Structural alignment of the extracellular region of wild-type and Alfa-tagged transferrin receptor (TFR). Wild type: PDB 1suv, while Alfa-tagged was generated by AlphaFold 3. See figure S1 for comparison of Alfa-tagged dimeric and monomeric TFR. **e**, Schematics showing the experimental design to test if transferrin receptor uptake is stimulated by the presence of transferrin. **f**, Quantification of transferrin receptor uptake for 15 or 30 minutes by flow cytometry using EndoNB in the presence of increasing concentrations of transferrin. (n = 4, each n represents the median fluorescence of 2000-8000 cells). See figure S1b for quantification of transferrin-AlexaFluor488 signals at each condition. **g**, Structural alignment of the transmembrane and extracellular region of wild-type and Alfa-tagged AXL. Both structures generated with AlphaFold 3. **h**, Quantification of AXL uptake by flow cytometry using EndoNB in the absence or presence of the AXL ligand GAS6 (1ug/ml). (n = 4, each n represents the median fluorescence of 2000-8000 cells). See figure S1c for comparison with the conditions with and without 3C. ns = non-significant, * p> 0.05, **** p> 0.0001. ANOVA with Tukey’s post hoc analysis.

Transferrin (TF) uptake via the transferrin receptor (TFR) is the most used tool to study clathrin-mediated endocytosis (CME). TFR is constitutively internalized, regardless of the presence of the TF ^24^. However, there are conflicting views in the literature on whether the presence of TF increases TFR internalization or not^25–29^. This type of question is a well-suited use case for EndoNB, so we generated a cell line Alfa-tagged for TFR (gene name TFRC) and performed our assays with increasing concentrations of TF (Fig. 2d,e and S1a). Our results detected no difference in internalized EndoNB signal among all conditions tested (Fig. 2f and S1b), in support of the view that TFR internalization rates are not affected by ligand presence or concentration.

The AXL receptor is a member of the receptor tyrosine kinase (RTK) family implicated in inflammatory processes such as immune-mediated cell clearance and blood clotting^30^. Crucially, AXL is involved in viral infections and cancers^31^ with drugs targeting this receptor being already in use or in late clinical development^32^. Akin to other RTK family members, AXL internalization is triggered by its ligand, growth arrest specific protein 6 (GAS6)^33^. To test if EndoNB can detect GAS6-stimulated AXL internalization, we generated a cell line Alfa-tagged (Fig. 2g) AXL and performed a time course internalization experiment. Our results showed that GAS6 led to an increase of AXL internalization which, under our experimental conditions, reached a plateau at 15 minutes (Fig. 2h and S1c), which is in line with what has been described previously^34^. Taken together, these results show that EndoNB is a simple and powerful tool to study the internalization behavior of receptors in the presence or absence of ligands and activity modulators.

### Applying EndoNB for direct comparison of receptor internalization

Next, we decided to further explore the differences between integrin β1 and integrin β5. It is fair to assume that the EndoNB probe will have similar fluorescence intensity and similar affinity to the Alfa-tag regardless of the receptor it targets. Therefore, we reasoned that absolute fluorescence values would allow a direct comparison of the internalization behavior of these receptors.

First, we generated a cell line with Alfa-tagged integrin β5. Here, we targeted the equivalent region we used for integrin β1 (Fig. 3a). To confirm the functionality of our tagged integrin β5, the staining of Alfa-tagged cells with EndoNB marked cellular adhesions which fully colocalized with an integrin αVβ5 staining (Fig. S2). Of note, Integrin αVβ5 is the only integrin pair formed by integrin β5. Next, we observed the internalization of integrin β5 by microscopy using an experimental design similar to our proof-of-principle experiment in figure 1e. This experiment revealed a very low levels of integrin β5 internalization, even after 3 h of incubation (Fig. S2).

**Figure 3.**
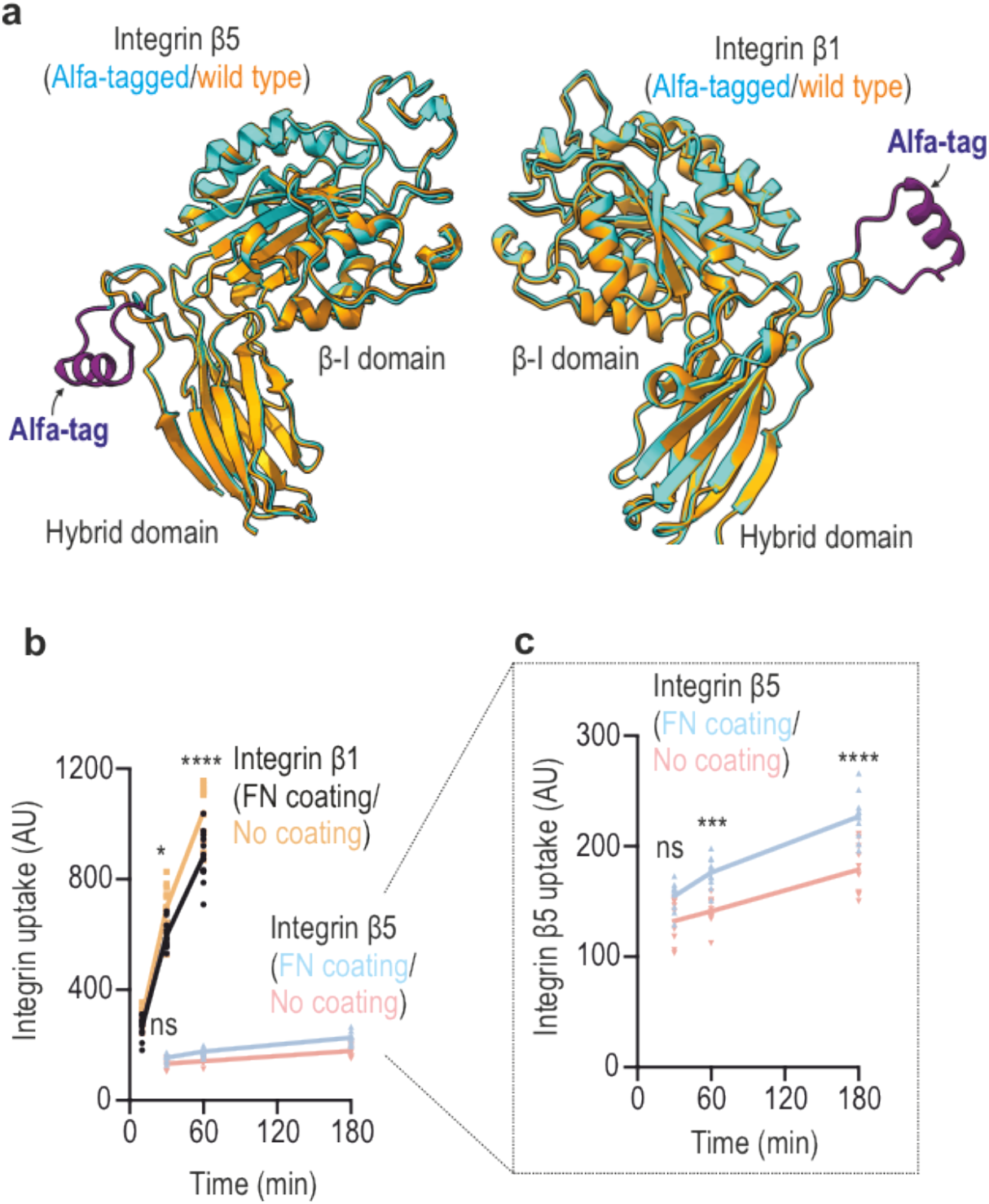
EndoNB to directly compare receptors. **a**, Structural alignment of the hybrid and β-I domains of wild-type and Alfa-tagged Integrin β1 and Integrin β5. **b**, Quantification of integrin β1 and integrin β5 uptake by flow cytometry using EndoNB comparing cells in uncoated plates vs fibronectin-coated plates. (n = 10, each n represents the median fluorescence of 2000-8000 cells). **c**, Graph inset showing a zoomed view of integrin β5 uptake. ns = non-significant, * p> 0.05, *** p> 0.001, **** p> 0.0001. ANOVA with Tukey’s post hoc analysis.

We then used integrin β5 and integrin β1 Alfa-tagged cells to directly compare their internalization rate in a time course experiment. Our results revealed that integrin β1 uptake was more than one order of magnitude faster when compared to integrin β5 (a slope 14 for β1 compared to a slope 0.3 for β5), (Fig. 3b). Integrin β5 is a unique integrin as it can localize to both focal adhesions (FAs) and reticular adhesions (RAs)^35,36^. We previously showed that this localization is controlled by the composition of the extracellular matrix (ECM), with uncoated dishes favoring integrin β5 on reticular adhesions (RAs) and fibronectin (FN) coated dishes favoring integrin β5 on focal adhesions (FAs)^36^. One key difference between integrin β5 in FAs and RAs is their turnover, with RAs being significantly more stable^35^. Confirming that this difference in stability is due to distinct endocytic rates, integrin β5 uptake was significantly faster in FN coated dishes (favoring integrin β5 on FAs) when compared to uncoated ones (favoring integrin β5 on RAs) (Fig. 3c). Interestingly, the uptake behavior of integrin β1 showed the opposite trend to integrin β5, with FN coating leading to slower uptake when compared to the uncoated condition (Fig. 3b). This effect is likely due to the fact that integrin β1 engagement with FN leads to its increased stabilization at focal and fibrillar adhesions. Hence, the use of a single and universal EndoNB probe allows the direct comparison of the internalization behavior of different surface proteins.

### Employing EndoNB to study poorly characterized surface proteins

One powerful application for EndoNB lies in its capacity to study unknown surface proteins. To showcase this potential, we generated a cell line with Alfa-tagged TMEM123 (also known as Porimin). TMEM123 is a highly glycosylated, mucin-like protein, linked to oncotic cell death and immune surveillance of colorectal cancers^37,38^. In a screen for CME cargoes, TMEM123 was identified as one of the top targets^39^. However, details on the function of this protein and its endocytic behavior are completely missing. The extracellular region of TMEM123 is predicted to be unstructured, so we added the Alfa tag to the N-terminal region following the signal peptide (Alanine 34) (Fig. 4a).

**Figure 4.**
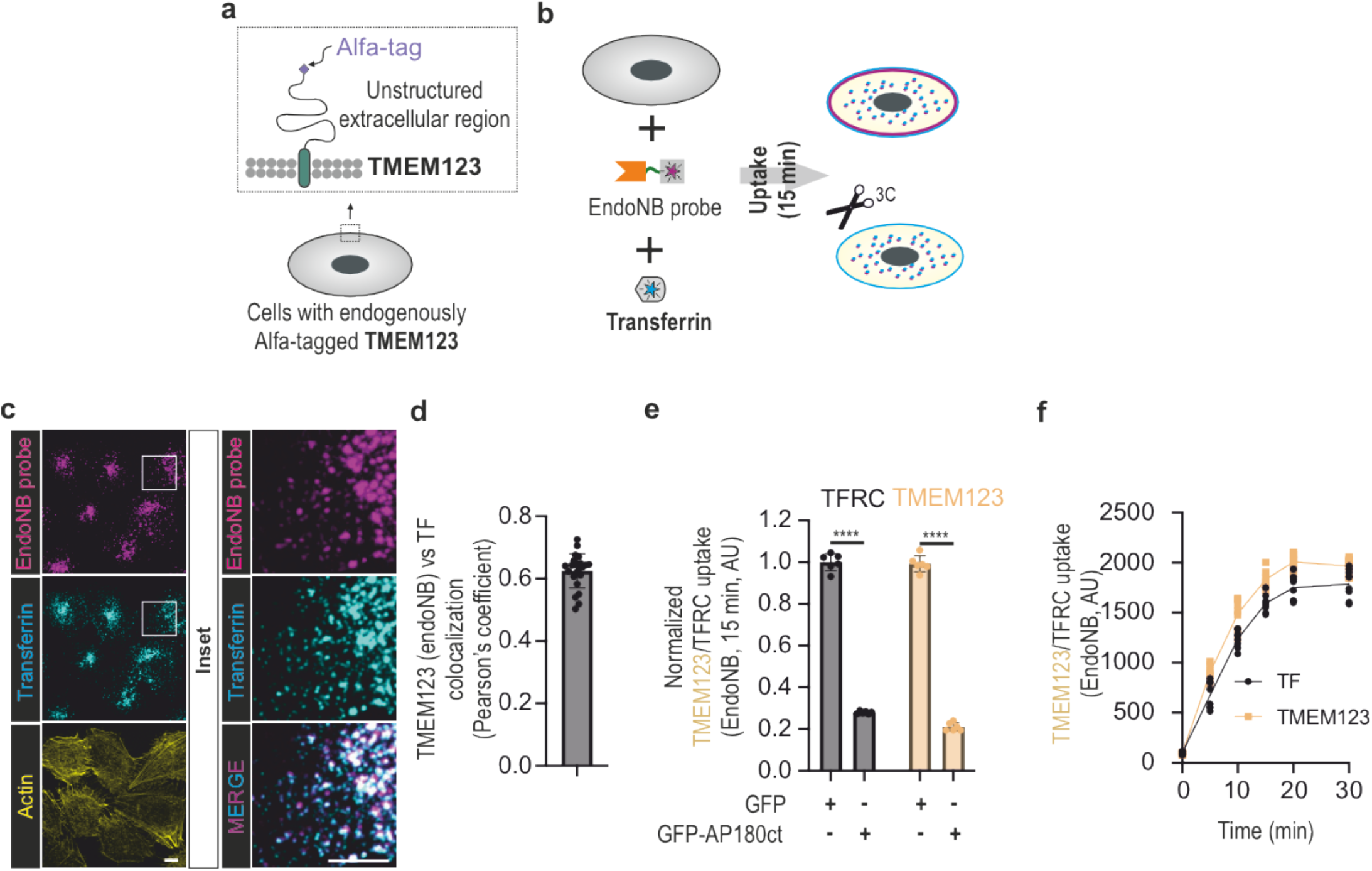
EndoNB to study poorly characterized surface proteins. **a**, Schematic showing the position of the Alfa-tag in the N-terminus (after the signal peptide) of the mucin-like surface protein TMEM123. The protein is predicted to be fully unstructured. **b**, Schematics showing the experimental setup to test the concomitant uptake of TMEM123 (using EndoNB) and transferrin. **c**, Images of cells after 15 minutes of TMEM123 (EndoNB) and Transferrin-AlexaFluor488 uptake. Images are maximum intensity projection of confocal images. See also figure S3 for the characterization of TMEM123 uptake using EndoNB. Scale bars = 10µm. **d**, Quantification of TMEM123 and transferrin colocalization. (n = 23 cells). **e**, Quantification of TMEM123 and TFR uptake by flow cytometry using EndoNB comparing cells transfected with the CME inhibitor AP180 c-terminus or GFP alone (n = 6, each n represents the median fluorescence of 2000-8000 cells). **f**, Time course uptake of TMEM123 and TFR uptake by flow cytometry using EndoNB (n = 7, each n represents the median fluorescence of 2000-8000 cells). It is important to note that this graph measures absolute uptake (not normalized), suggesting that the endocytic behavior of these two cargoes is identical. ns = non-significant, **** p> 0.0001. ANOVA with Tukey’s post hoc analysis.

Our initial experiments revealed that TMEM123 shows robust endocytosis within 15 minutes (Fig. S3). Due to its apparent kinetic similarity with TFR, we decided to combine TMEM123 uptake (EndoNB) with TF uptake (Fig. 4b). Confirming that these two cargoes share a common endocytic fate, a significant amount of TMEM123 colocalized to endosomes containing TF (Fig. 4c,d and S3). Moreover, both cargoes showed similar reduction in uptake in cells expressing a CME inhibitor (The C-terminus of the AP180 protein, a potent dominant-negative inhibitor of CME^40^) (Fig. 4e). Given the similarities between these cargoes, we decided to compare their uptake rates side-by-side using TMEM123 and TFR Alfa-tagged cells. To our surprise, their uptake profile was remarkably similar, with a steady rise for 15-20 minutes followed by a plateau around 20 minutes (Fig. 4f). Thus, EndoNB can be used to study novel surface proteins for which tools are currently unavailable.

## Discussion

Here we show that EndoNB is a robust and flexible assay capable of measuring the internalization of virtually any surface protein using different readout modalities and for multiple purposes. Our assay removes the limitations imposed by the rarity of good antibodies^41^ or engineered ligands, thereby enabling researchers to study the biology of any surface protein with ease. Moreover, we foresee that the flexibility of EndoNB allows refinements on the breadth and specificity of known endocytic pathways and unlocks the discovery of new ones.

Using EndoNB, we were able to: (i) study the endocytosis of novel surface proteins; (ii) reproduce the endocytic behavior, (iii) directly compare the internalization and, (iv) address new questions on established receptors. For example, we could show that a widely used inhibiting antibody for integrin β1^20^ doubles the internalization of the total population of this protein. Another interesting finding was the remarkable similarity between the endocytic behavior of TMEM123, an abundant but poorly studied surface protein^37,38^, and the Transferrin receptor TFR, the canonical marker for CME. We found that the absolute amount of internalization, uptake kinetics, endosomal localization and response to CME inhibitors for these proteins were almost identical. Given that their expression is in the same range in the cell line we used, U2Os (according to The Human Protein Atlas^42^), these similarities suggest that constitutive CME has a specific kinetic signature.

A key limitation of EndoNB lies in the expression level of the target surface protein in the cells of interest. In our experience, mRNA expression levels above 30-50 nTPM (normalized transcripts per million, data from The Human Protein Atlas^42^) are required to yield enough signal for reliable detection of surface and internalized signals. It is likely that strategies to increase the signal on the fluorescent moiety of the EndoNB probe could solve this issue.

Looking at the future, it is tempting to speculate that the basic principle of EndoNB could leverage latest developments in the generation of synthetic protein binders^43,44^ to enable our assay in unmodified cells. We envisage that by replacing the nanobody Alfa from EndoNB by different synthetic binders targeting various surface proteins, and labelling them with different fluorophores, one could follow the internalization behavior of many surface proteins simultaneously in unedited primary cells or even *in vivo*.

The tale of the blind men and the elephant illustrates how each man, by touching only a part of the animal (e.g., trunk, tail or tusk), claims to have a knowledge of the whole. To a certain extent, our knowledge on endocytic and trafficking pathways suffers from the same blindness, with most of our knowledge relying on a very limited number of experimentally traceable proteins. The approach we present here removes many of these previous limitations and has the potential to unlock biological insights of thousands of new proteins in both physiological and disease contexts.

## Acknowledgements

We would like to thank the Lavis lab and Open chemistry team (Janelia) for the gift of SNAP-JF substrates. We would like to thank the HiLIFE Light Microscopy Unit (LMU) and the HiLIFE Flow Cytometry unit for technical assistance. The authors wish to acknowledge CSC – IT Center for Science, Finland, for computational resources and the Bioimaging and optic platform (BIOP-Ecole Polytechnique fédérale de Lausanne) for the easy-to-use ImageJ implementation of Cellpose. LA-S is supported by HiLIFE, the Academy of Finland (Research Fellow) and Sigrid Juselius Foundation (Young PI grant). J.K. was supported by a postdoctoral research grant by the Estonian research council (PUTJD1114). We would like to thank Pekka Lappalainen (UH) and Brendan Battersby (UH) for the critical and kind reading of our manuscript. The authors declare no competing financial interests.

## Author contributions

*Investigation* – An-Sofie Lenaerts, Jekaterina Krishtal, Tai Arima, Neza Leskovar, Narjes Zeinoddin, Leonardo Almeida-Souza

*Methodology* - An-Sofie Lenaerts, Jekaterina Krishtal, Tai Arima, Leonardo Almeida-Souza.

*Conceptualization, Supervision, Formal analysis, Visualization, Funding acquisition* - Leonardo Almeida-Souza.

*Writing* - Leonardo Almeida-Souza with input from all authors.

## Materials and Methods

### Cell culture and reagents

U2Os cells and all its engineered derivatives were cultured in MEM supplemented with 10% fetal bovine serum (FBS) (Gibco) and penicillin-streptomycin (100 U/ml, Thermo Fisher Scientific). HEK293T Flp-in cells were grown in DMEM media supplemented with 10% fetal bovine serum (FBS) (Gibco), penicillin-streptomycin (100 U/ml, Thermo Scientific), 5µg/ml blasticicin and 100µg/ml zeocin (Host cell line) or 100µg/ml hygromycin (stable cell line).

SNAP-JF646 substrate was a kind gift by Luke Lavis via the open chemistry team (jJanelia); Transferrin-Alexa Fluor 488 (Thermo Fisher scientific, #T13342); Phalloidin-iFluor 488 (AAT Bioquest, #23115); GAS6 recombinant protein (MedChemExpress via Nordic Biosite HY-P700724-20).

The following primary antibodies were used: anti-human integrin β1 activating antibody 12G10 (Novus bio, NB100-63255), anti-human integrin β1 inactivating antibody mAb13 (BD, #552828), anti-human integrin αVβ3 (Invitrogen, #11-0519-42), anti-human integrin αVβ5 clone 15F11 (MAB2019Z; Millipore).

Transient transfections were carried out with PEI MAX transfection reagent (Polysciences, 24765-1).

### Constructs

gRNAs were ordered as primers (from IDT) and cloned into pSpCas9(BB)-2A-Puro (PX459) V2.0 (gift from Feng Zhang, Addgene #62988) using BbsI sites and confirmed by Sanger sequencing.

The EndoNB probe was synthesized by IDT as gene blocks. For E.coli expression, this gene block was cloned into p7XH3 (Addgene, #47064) using fx cloning. For mammalian expression, the gene block containing the igKVII (immunoglobulin kappa variable cluster) signal peptide (MDMRVPAQLLGLLLLWLRGARC) was cloned using gateway into the flp-in system-ompatible vector MAC-tag-C45. Please note that the mac-tag itself was not expressed in our constructs.

The CME inhibitor AP180 c-terminal fragment (amino acids 516-898) cDNA, from rat origin, was described previously46. This construct was cloned into Gateway compatible pCI vectors, containing an N-terminal monomeric EGFP using the Gateway system.

### Generation of Alfa-tagged cells

Sites for Alfa-tag insertion were defined on previous literature reports of internal tagging sites, structural and functional data, AlphaFold 2^17^ or AlphaFold 3^16^ predictions and combinations thereof. In general terms, we aimed to add Alfa-tags to unstructured loops, at positions away from dimerization interfaces, ligand binding sites and glycosylation sites.

For each surface protein, we designed 2-3 guide RNAs (gRNA) sequences using the Wellcome Sanger Institute online genome editing tool (https://wge.stemcell.sanger.ac.uk/). gRNAs were cloned into pSpCas9(BB)-2A-Puro (PX459) V2.0 as described above.

Single strand oligodeoxynucleotides (ssODN) were used as templates for homologous recombination. They were designed to contain 60-75 homology arms surrounding the Alfa-tag sequence (CCCAGCAGGCTGGAGGAAGAGCTGAGGCGCAGACTGACCGAGCCC) and flanked by a glycine-serine flexible linker.

70-80% confluent 6-well plates of U2Os cells were transfected with 10 µg PEI (1 µg/ml), 750 ng of plasmid and 12.5 pmol ODNs. Two days after transfection cells were treated for 48h with puromycin (1 µg /ml) to enrich for successfully transfected cells. The success for each gRNA was evaluated around 1-2 weeks of recovery after transfection, when cells were treated with the EndoNB probe (or a labeled nanobody anti-alfa, Nanotag, #N1502) for 1h at 37°C and FACS analyzed for the presence of positive cells. The most effective gRNA, judged by the percentage of fluorescent cells by FACS, was used for single clone sorting, genotyping and confirmation by microscopy. The region of integration was sequence confirmed for all clones used in experiments. All cells we used in experiments were heterozygous for the Alfa-tag insertion.

A full list of gRNA, ODNs and genotyping primers for genome editing is shown below:

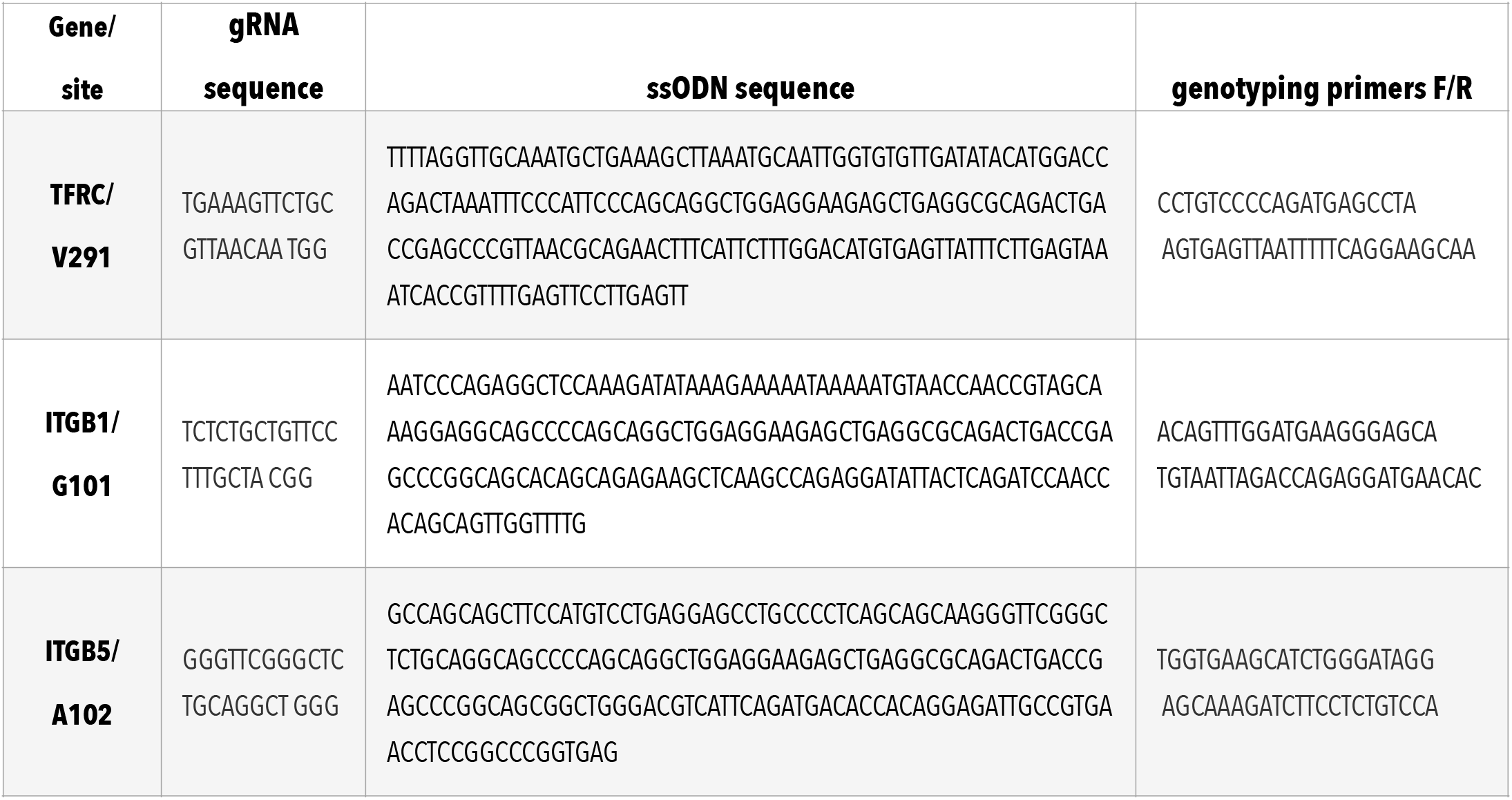
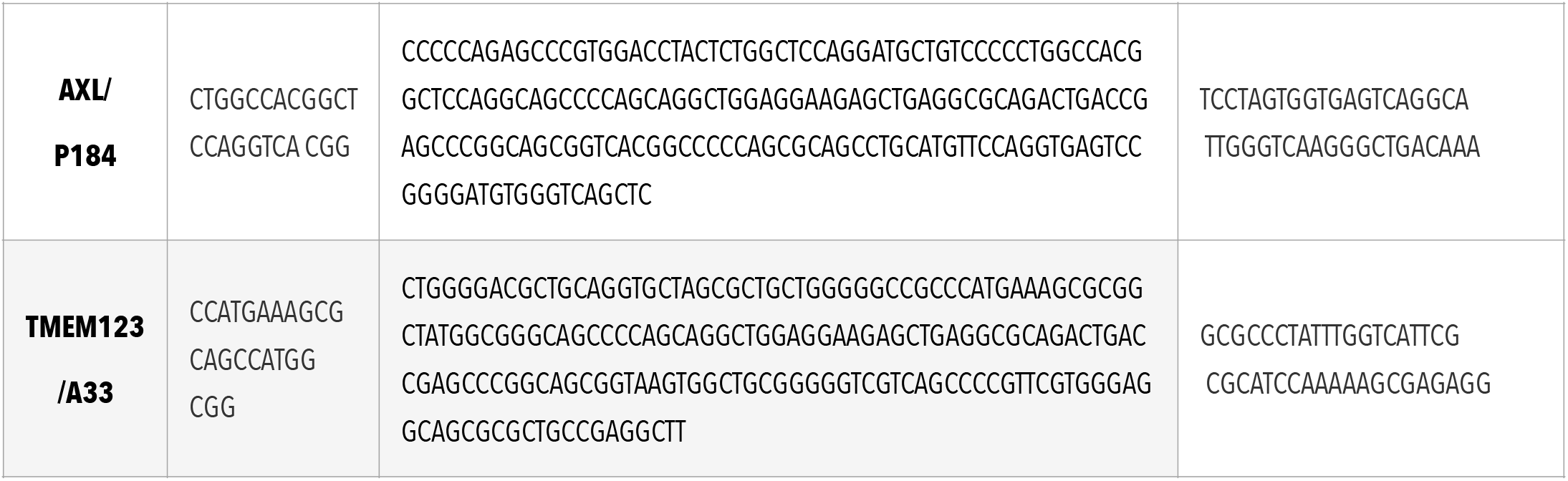

### Production of the EndoNB probe

Most of the optimization and preliminary data collected in this project were performed using the EndoNB probe purified from *E. coli*. At a later point, we optimized the production of EndoNB from mammalian cells and the results were similar when using either source of probe. Even though most of the results presented here were performed with the mammalian produced probe, we decided to include both production protocols to accommodate a wide range of laboratory setups.

#### E.coli

In E. coli, the EndoNB probe could not be expressed in the soluble fraction and had to be purified from pellets and refolded. The protein was expressed in Rosetta (DE3) *Escherichia coli* overnight at 18°C. Cells were harvested via centrifugation at 4350rcf and resuspended in 1:1 (vol/weight) of lysis buffer (50 mM Tris pH 8, 500 mM NaCl, 30 mM imidazole). Cells were then treated with 1mg/ml of lysozyme for 15 min at room temperature and lysed by sonication using a probe sonicator (VC-750 Vibra-Cell Ultrasonic Liquid Processor, Fischer Scientific) and spun at 21,000 rcf for 30 minutes at 4°C. The insoluble fraction was retained and washed four to five times using 1% Triton (v/v) in lysis buffer (1 mL of this wash buffer per gram of pellet) to remove remaining lipid components from the pellet. Proteins were then solubilized with 6 M guanidine followed by vigorous mixing using a spatula and 30-minute incubation at 37° C. This solution was then centrifuged at 21,000 rcf for 10 minutes, and the supernatant (containing denatured, resolubilized EndoNb) was retained, while the pellet was discarded. The proteins, still in guanidine, were passed through a HisTrap HP column (Cytvia, #17524802), extensively washed with 6 M guanidine solution and eluted with 6 M guanidine supplemented with 300 mM imidazole. For refolding, a refolding buffer was prepared with the following components: 50 mM Tris-HCl pH 8, 5 mM EDTA, 500 mM NaCl, 500 mM Arginine, 3 mM glutathione, and 0.3 mM glutathione disulfide. The refolding buffer solution was created at a 20:1 volumetric ratio to the His-tag purification elution containing denatured EndoNb and set to stir rapidly. The His-tag purification elution was then added dropwise and allowed to continue stirring for one hour after the complete volume had entered the refolding buffer solution. All steps were performed at 4° C. Following refolding, the protein was centrifuged 21,000 rcf for 30 minutes to remove insoluble aggregates and concentrated to a volume of approximately 1 mL using Amicon® Ultra 30K Centrifugal Filters (Merck). The concentrated refolded protein was incubated overnight at 4°C with a 2-molar fold excess of SNAP-JF646 substrate and further purified using a Superdex 200 (GE healthcare) size exclusion column using gel filtration buffer (150 mM NaCl, 20 mM HEPES pH 7.5, 500 μM TCEP). The final protein was concentrated to 0.5 mg/ml and frozen in 20µl aliquots.

#### Mammalian cells

First, we generated cells stably expressing EndoNB using EndoNB cloned in the MAC-tag-C vector and Flp-in T-Rex HEK293T cells (ThermoFisher Scientific) according to manufacturer instructions. For expression, cells were grown to 70% confluence in one 1750cm^2^ Hyperstack flask (Corning, #CLS10031) and media was changed to DMEM media supplemented with 2% fetal bovine serum (FBS) (Gibco), 5µg/ml Blasticicin, 100µg/ml hygromycin and 1µg/ml tetracycline (for EndoNB induction). Media was collected and replaced every 2-3 days over a 10-day period. Media was concentrated 100-500-fold using Amicon® Ultra 30K Centrifugal Filters (Merck), diluted in binding buffer (50 mM Tris pH 8, 100 mM NaCl, 30 mM imidazole, final concentration), bound to HiTrap Q-Sepharose FF columns (3×5ml in tandem, Cytiva, #17515601) and eluted using a 100-500 NaCl gradient. Fractions containing EndoNB were supplemented to ∼500mM NaCl and injected on a HisTrap HP column (Cytiva, #17524802), extensively washed with 50 mM Tris pH 8, 500 mM NaCl, 30 mM imidazole, and eluted with 50 mM Tris pH 8, 500 mM NaCl, 300 mM imidazole. The eluate was concentrated using Amicon® Ultra 30K Centrifugal Filters (Merck), incubated overnight at 4°C with a 2-molar fold excess of SNAP-JF646 substrate and further purified in a Superdex 200 (Cytiva) size exclusion column using gel filtration buffer (150 mM NaCl, 20 mM HEPES pH 7.5, 500 μM TCEP). The final protein was concentrated to 0,5 mg/ml and frozen in 20µl aliquots.

### Uptake assays

#### Flow cytometry

25000 Alfa-tagged cells/well were seeded in 96 well plates. On the next day, cells were incubated with EndoNB probe (1 µg/ml) and/or transferrin-AlexaFluor 488 (10 µg/ml, unless stated otherwise) at 37°C for the time described in the graphs for each experiment. After the incubation, cells were placed on ice to stop internalization and washed with ice-cold PBS. To remove surface fluorescence, cells were incubated with HRV 3C protease (2 µg/ml in PBS) for 1h at 4°C. When described, a control condition without 3C (only PBS) was also incubated for 1h at 4°C. After 3C treatment, cells were washed with PBS twice, detached using Accutase (Sigma, A6964), transferred to 96 well V-bottom plates and analyzed using an LSR fortessa (BD biosciences) equipped with a high throughput sampler (HTS). We only used conditions with a minimum of 2000 valid events.

#### Microscopy

50000 Alfa-tagged cells/well were seeded in 24 well plates containing 13mm diameter coverslips. On the next day, cells were incubated with EndoNB probe (1 µg/ml) and /or transferrin-AlexaFluor 488 (10 µg/ml) at 37°C for the time described in the graphs for each experiment. After incubation, cells were placed on ice to stop internalization and washed with ice-cold PBS. To remove surface fluorescence, cells were incubated with HRV 3C protease (2 µg/ml in PBS) for 1h at 4°C. When described, a control condition without 3C (only PBS) was also incubated for 1h at 4°C. After 3C treatment, cells were washed with PBS twice, fixed with 4% paraformaldehyde in PBS for 15 min in a +37°C incubator, washed with PBS followed by staining with Phalloidin-AlexaFluor568 (30min, room temperature, 66nM or 1000x dilution of 400x stock). For integrin αVβ5 antibody staining additional steps after fixation were included: blocking with 1% BSA-PBS (30min, room temperature), primary antibody staining (1h, room temperature, 1:600), washing with PBS, secondary antibodies conjugated with fluorophores were applied together with Phalloidin AlexaFluor568 (30min, room temperature, 1:1000). Coverslips were then mounted using Immu-Mount™ (epredia) overnight at 4°C. Images were acquired using a 40x (HC PL APO 1.25NA) objective in a Leica Stellaris SP8 equipped with a UV and a White light laser (440-790).

#### Cell specific protocol modifications and concentration of treatments

For experiments using transferrin (or transferrin receptor) uptake, we serum starved the cells for 30 minutes before adding transferrin or the EndoNB probe. When specified, plates were coated with 10µg/ml of Fibronectin (Merck, 341631) overnight at 37°C. For Integrin β5 cells we used 15000/30000 cells/well in 96/24 well plates. This was due to the fact that we have previously observed that high confluence masks the effects of fibronectin on Integrin β5 behaviour^36^. Concentration of treatments: 12G10 = 250ng/ml; Mab13 = 0,15ng/ml; αvb3 = 0,1ng/ml; Gas6 =1ug/ml.

### Image Analysis

All images were analyzed using ImageJ^47^. To quantify uptake, we used an ImageJ implementation of Cellpose^48^ to automatically segment cells. Cells were segmented using the cyto2 algorithm of Cellpose^48^ using the phalloidin channel as input. Uptake was measured for each cell as fluorescence integrated density. To measure colocalization of transferrin and TMEM123, we used the ImageJ plugin Coloc 2.

### Statistical analysis

All experiments were repeated at least 3 times with 2-4 replicates. We opted to present primarily absolute fluorescence values for uptake and, when possible, we pooled different experiments into the same graph. In other cases, when absolute values were different, we opted to show a representative experiment with a minimum of 4 replicates. Nevertheless, we only presented results whose trend between different experiments was consistent. For multiple comparisons one-way ANOVA was performed followed by Tukey’s multiple comparison. All graphs and statistical calculations were performed with GraphPad Prism 9.

## Supplementary Figures

**Figure S1.**
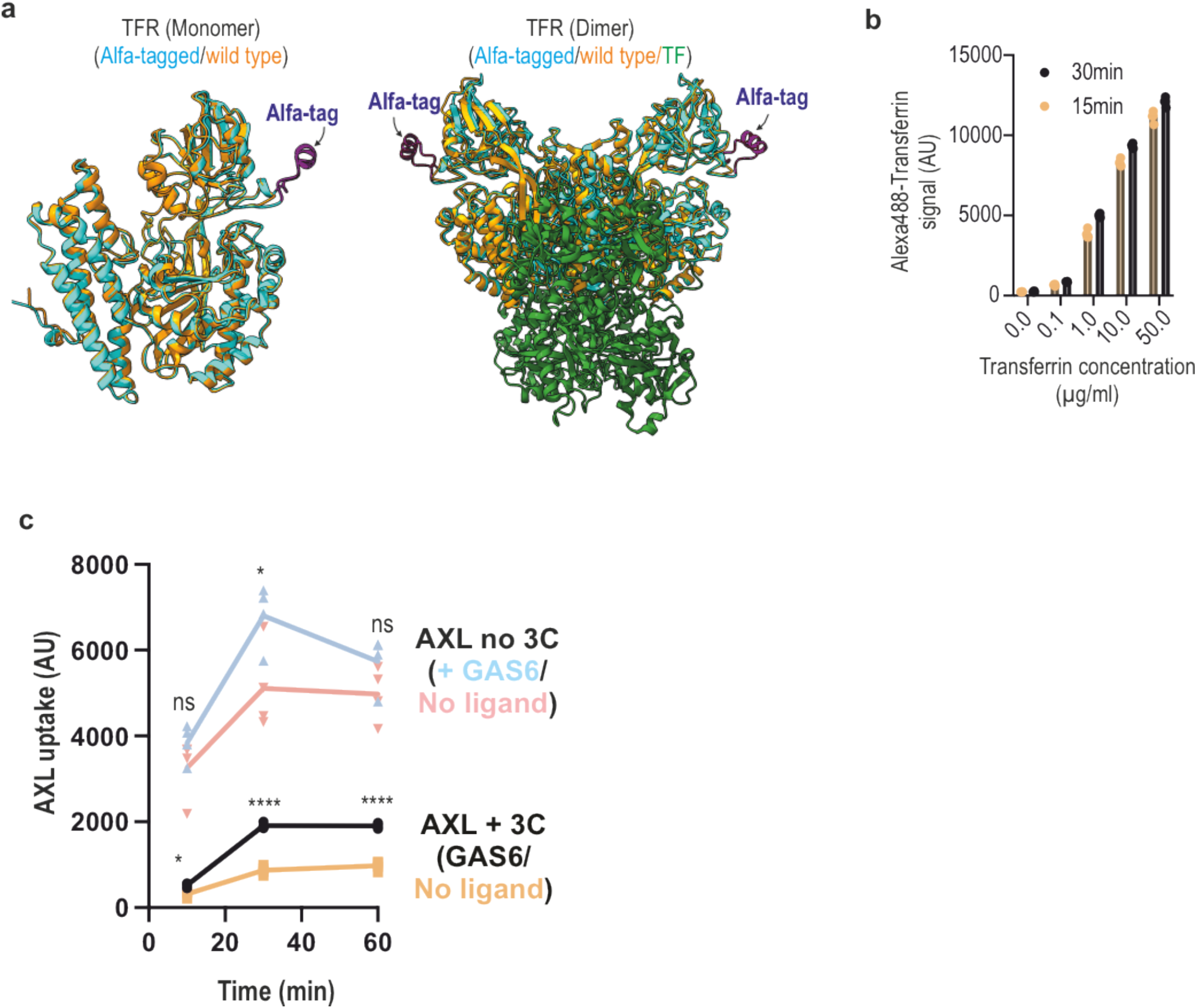
**a**, Structural alignment of the monomeric and dimeric extracellular region of wild-type and Alfa-tagged transferrin receptor (TFR, PDB). Wild type: PDB 1suv, while Alfa-tagged was generated by AlphaFold 3. **b**, Quantification of transferrin (AlexaFluor488) in each condition measured in figure 2f. **c**, Quantification of AXL uptake by flow cytometry using EndoNB in the absence or presence of the AXL ligand GAS6 (1µg/ml) and also in the presence or not of 3C. (n = 4, each n represents the median fluorescence of 2000-8000 cells). ns = non-significant, ** p> 0.01, *** p> 0.001, **** p> 0.0001. ANOVA with Tukey’s post hoc analysis.

**Figure S2.**
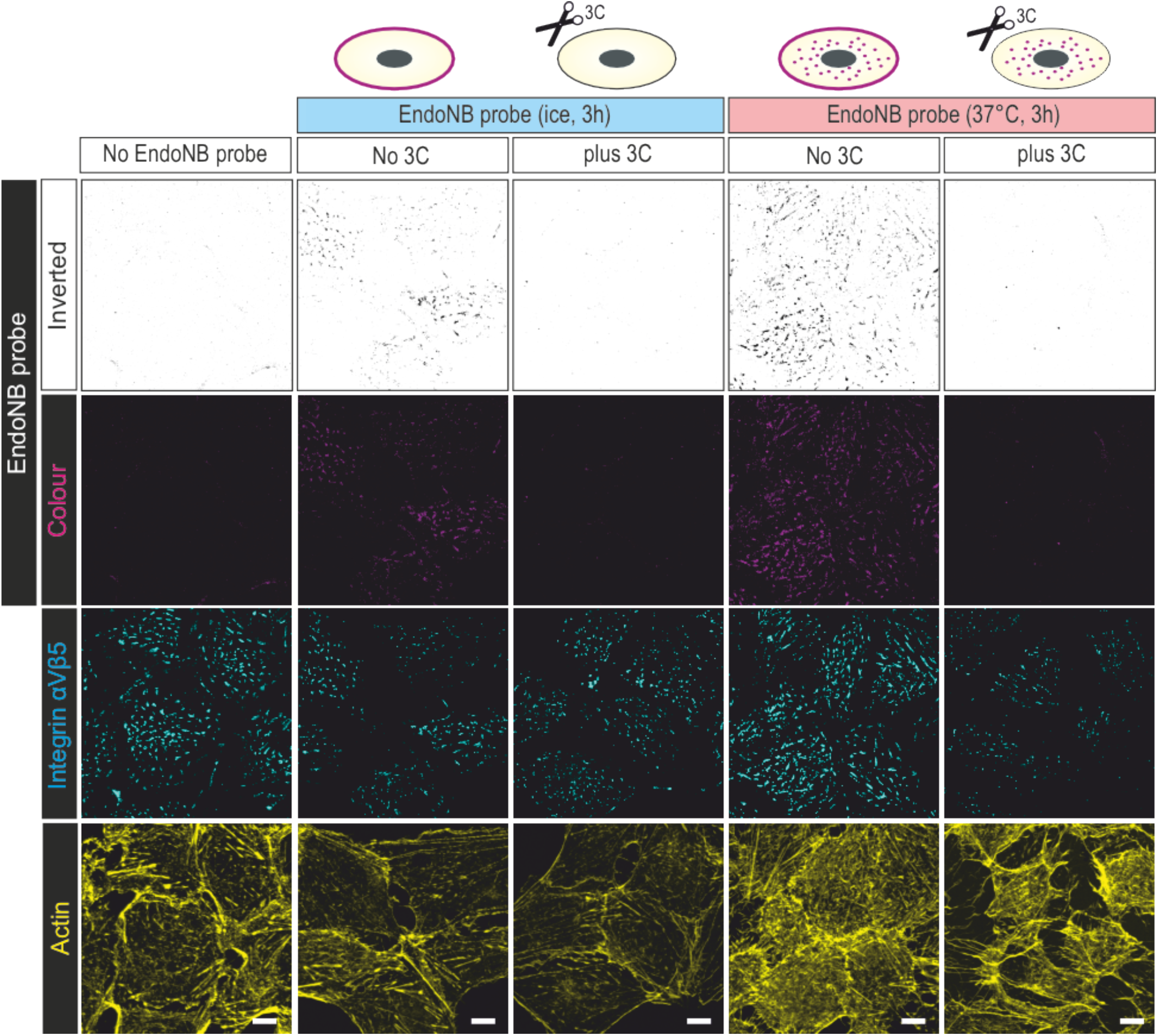
Characterization of EndoNB uptake in Alfa-tagged integrin β5 cells. Images are maximum intensity projection of confocal images. Note that Alfa-tagged integrin β5 still fully colocalizes to cell adhesions (as judged by the αVβ5 staining). Note also the slow uptake of integrin β5. Scale bar = 10µm.

**Figure S3.**
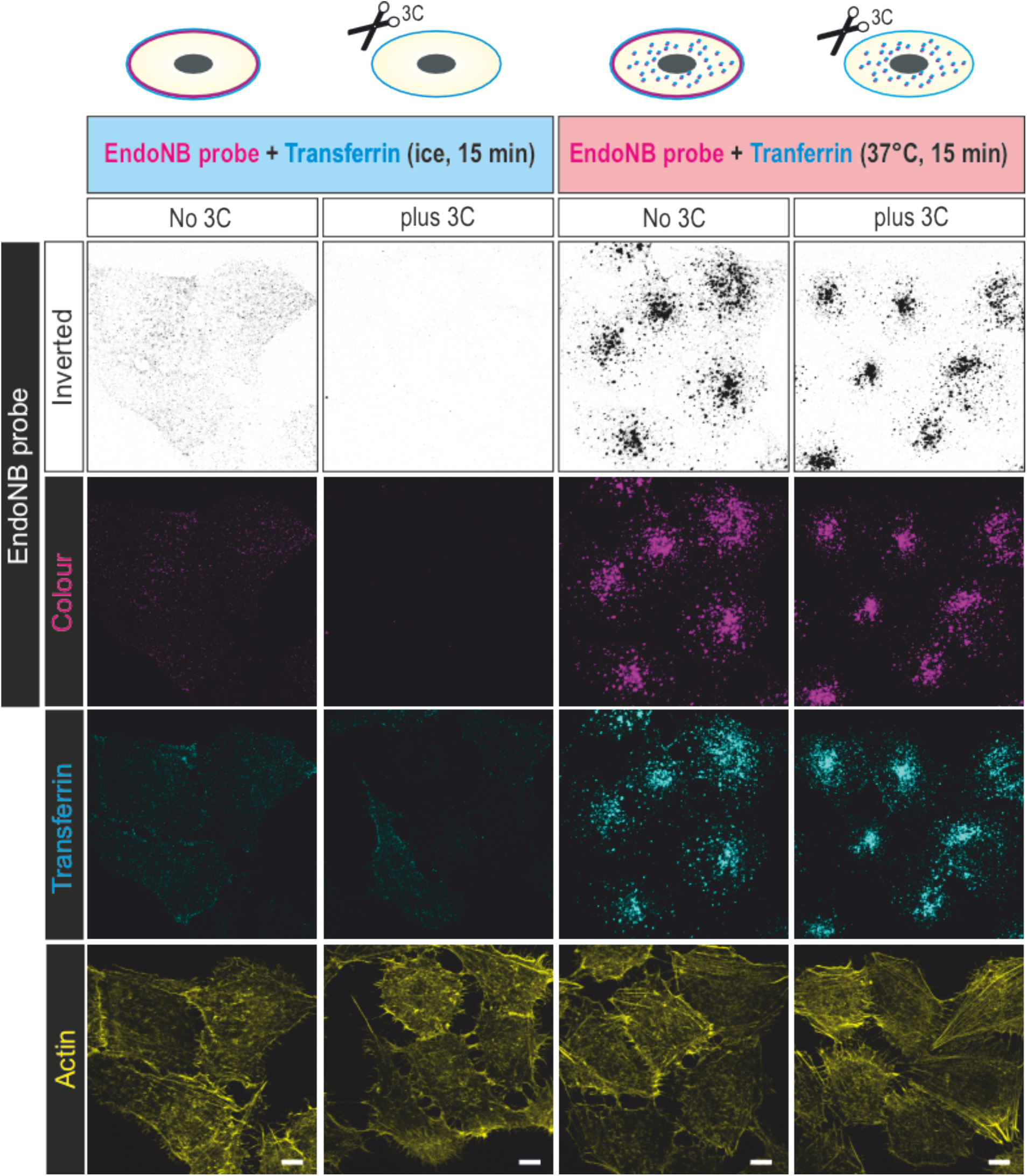
Characterization of EndoNB uptake in Alfa-tagged TMEM123 cells. Images are maximum intensity projection of confocal images. Scale bar = 10µm.

## Notes

### Competing Interest Statement

The authors have declared no competing interest.

